# *Δ*-*d*_*N*_/*d*_*S*_: A New Criteria to Distinguish among Different Selection Modes in Gene Evolution

**DOI:** 10.1101/2020.02.21.960450

**Authors:** Xun Gu

## Abstract

One of the most widely-used measures for protein evolution is the ratio of nonsynonymous distance (*d_N_*) to synonymous distance (*d_S_*). Under the assumption that synonymous substitutions in the coding region are selectively neutral, the *d_N_*/*d_S_* ratio can be used to test the adaptive evolution if *d_N_*/*d_S_*>1 statistically significantly. However, due to selective constraints imposed on amino acid sites, most encoding genes demonstrate *d_N_*/*d_S_*<1. As a result, *d_N_*/*d_S_* of a gene is less than 1, even some sites may have experienced positive selections. In this paper, we develop a new criterion, called *Δ-d_N_*/*d_S_*, for positive selection testing by introducing an index *H*, which is a relative measure of rate variation among sites. Under the context of strong purifying selection at some amino acid sites, our model predicts *d_N_*/*d_S_*=1-*H* for the neutral evolution, *d_N_*/*d_S_*<1-*H* for the nearly-neutral selection, and *d_N_*/*d_S_*>1-*H* for the adaptive evolution. The potential of this new method for resolving the neutral-adaptive debates has been illustrated by case studies. For over 4000 vertebrate genes, virtually all of them showed *d_N_*/*d_S_*<1-*H*, indicating the dominant role of the nearly-neutral selection in molecular evolution. Moreover, we calculated the *d_N_*/*d_S_* ratio for cancer somatic mutations of a human gene, specifically denoted by *C_N_*/*C_S_*. For over 4000 human genes in cancer genomics, about 55% of genes showed 1-*H*<*C_N_*/*C_S_*<1, about 45% of genes showed *C_N_*/*C_S_*<1, whereas less than 1% of genes showed *C_N_/C_S_*<1-*H*. Together our analysis suggested driver mutations, i.e., those initiate and facilitate carcinogenesis, confer a selective advantage on cancer cells, leading to *C_N_*/*C_S_*>1 (strong positive selection) or 1-*H*<*C_N_*/*C_S_*<1 (weak positive selection, combined with strong purifying selection), whereas nearly neutral selection due to reduced effective clonal size is highly unlikely in cancer evolution.

## Introduction

Since the proposal of neutral theory (Kimura 1968; King and Jukes 1969) as well as the more general nearly-neutral theory (Ohta 1973; Kimura 1983), the debate between selectionists and neutralists has continued to the era of genome sciences (Lynch 2007; Hahn 2008; Fay et al. 2011; Lynch et al. 2016; Kern and Hahn 2018; Jensen et al. 2019). This is partly because detection of positive selection at the genome level remains a challenge in germline mutations (Bustamante et al. 2005; Sabeti et al. 2006) or in somatic mutations in cancers (Chen et al. 2019a). One of most widely-used measure for protein sequence evolution is the ratio of nonsynonymous distance (*d_N_*) to synonymous distance (*d_S_*) (Li 1997). The *d_N_*/*d_S_* ratio has been widely used to test the underlying selection mode. Under the assumption that synonymous substitutions in the coding region are selectively neutral, the *d_N_*/*d_S_* ratio is a conventional estimate for the ratio of mean evolutionary 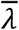 to the mutation rate (*v*) (Kimura 1983). It appears that the ratio *d_N_*/*d_S_*=1 means a strict neutral evolution in nonsynonymous substitutions; *d_N_*/*d_S_*>1 for positive selection and *d_N_*/*d_S_*<1 for negative selection.

However, the null hypothesis of *d_N_*/*d_S_*=1 (strictly neutrality) can be confounded by some amino acid sites subject to very strong purifying selection (lethal mutations) (Ohta 1992). Therefore, without knowing the proportion of lethal mutations to make some corrections, the *d_N_*/*d_S_* ratio of a gene could be less than one even when some sites are indeed subject to positive selection. In other words, the application of *d_N_*/*d_S_*>1 as a criterion to detect positive selection becomes less powerful. Indeed, the observation of *d_N_*/*d_S_*<1 can be explained by either the existence of essential sites plus some sites under (weak) positive selection, or the existence of essential sites in the dominance of strictly neutral evolution, or the nearly nearly-neutral evolution. In this paper, we propose a new criterion to distinguish between these possibilities.

## New Methods

### Evolutionary rate of an encoding-gene

In the theory of molecular evolution, the evolutionary rate (λ) of a nucleotide is determined by three levels of genetic processes: mutation at the individual level, polymorphism at the population level, and fixation at the species level (Kimura 1983). Let *v* be the mutation rate, *s* the coefficient of selection and *N_e_* the effective population size. It has been showed that the evolutionary rate is given by

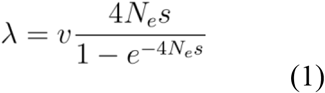

Eq.(1) has provided a solid foundation for resolving the debate between adaptive evolution, neutral evolution, and nearly-neutral evolution: it predicts λ/*v*>1 for adaptive evolution (*s*>0), λ/*v*=1 for neutral evolution (*s*=0), or λ/*v*<1 for deleterious evolution (*s*<0), respectively. Usually, *S*=*4N*_*e*_*s* is called the selection intensity that uniquely determines the rate-mutation ratio λ/*v*.

In practice, the ratio of nonsynonymous distance (*d_N_*) to synonymous distance (*d_S_*) is widely used in molecular evolution. Under the assumption that synonymous substitutions are selectively neutral, the *d_N_*/*d_S_* ratio can be used as a proxy of the λ/*v* ratio. Based on the *d_N_*/*d_S_* ratio, numerous statistical methods (Li et al. 1985; Nei and Gojobori 1986; Li 1993; Ina 1995; Cameron 1995; Yang 1997; Yang 2006) were developed to test the neutrality: rejection of the null hypothesis *d_N_*/*d_S_*=1 indicates a positive selection if *d_N_*/*d_S_*>1, or a negative selection if *d_N_*/*d_S_*<1. However, the *d_N_*/*d_S_* ratio is the mean estimated from the nucleotide sites of an encoding gene. Because each site may have different selection intensity (*S*), without loss of generality we assume that *S* varies among sites according to a distribution Φ(*S*). It follows that the *d_N_*/*d_S_* ratio is actually expected to be

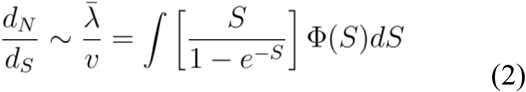

While the criteria for detecting different evolutionary modes remain the same, we have realized that the interpretation of *d_N_*/*d_S_* ratio test depends on the structure of *Φ*(*S*). For instance, with a few exceptions, it has been observed that *d_N_*/*d_S_*<1 holds for encoding genes of all organisms, indicating the role of selective constraints imposed on the protein sequence. There are two common interpretations of *d_N_*/*d_S_*<1. One is the nearly-neutral model, that is, the range of *S* in *Φ*(*S*) is negatively continuous, with a range of (−∞, 0); the other one is the neutral-lethal model, that is, *Φ*(*S*) follow a simple two-state distribution, *S*=0 (strictly neutral) and *S*=−∞ (lethal).

### The *Δ*-criterion on protein sequence evolution

It appears that additional information is required to develop a new *d_N_*/*d_S_*-based method that can distinguish between more sophisticated selection modes. Noting that the conventional *d_N_*/*d_S_* tests were actually focused on the mean rate-mutation 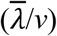 ratio, we consider the second moment of the evolutionary rate, that is,

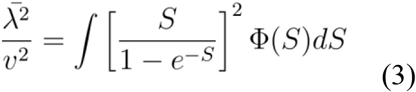

Next, we introduce a new quantity *Δ*, the difference between the second-moment and the mean of rate-mutation ratio. From Eq.(2) and Eq.(3), *Δ* can be written as follows

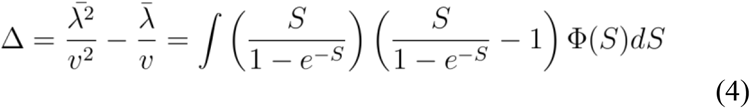

As shown below, the sign of *Δ* provides some insights for testing different selection modes.

#### Δ=0 under the neutral-lethal selection mode

Under the classical neutral model (Kimura 1968; 1983), the distribution *Φ*(*S*) can be constructed as follows: all mutations are classified into two categories: strictly neutral mutations (*S*=0 with a probability of 1−*f_L_*) or lethal mutations (*S*=−∞ with the probability of *f_L_*). Since the evolutionary rate is λ=*v* (mutation rate) for strictly neutral mutations and λ=*0* for lethal mutations, respectively, the first (mean) and the second moments of evolutionary rate of a gene under this model is simply given by

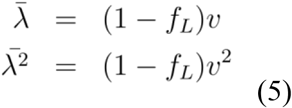

respectively. We thus conclude that, under the neutral-lethal selection mode, we have *Δ=0,* regardless of the proportion of lethal mutations.

#### Δ>0 under the positive selection with neutral-lethal mode

Suppose that all mutations of a gene are classified into three categories: adaptive mutations (*S*>0 with a probability of *f_A_*), lethal mutations (*S*=−∞ with a probability of *f_L_*), and neutral mutations (*S*=0 with a probability of 1−*f_A_*−*f_L_*). Moreover, the (positive) selection intensity of adaptive mutations follows a distribution denoted by *Φ^+^*(*S*). It follows that *Δ* can be written as follows

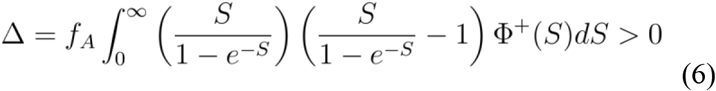

which is always larger than 0, regardless of the existence of lethal or neutral mutations.

#### Δ<0 under negative selection with nearly-neutral mode with lethal mutations

Under the nearly-neutral model with lethal mutations, all mutations of a gene are classified into three categories: nearly-neutral mutations (*S*<0 with a probability of 1−*f_L_*−*f_0_*), lethal mutations (*S*=−∞ with a probability of *f_L_*), and neutral mutations (*S*=0 with a probability of *f_0_*). Moreover, the (negative) selection intensity of nearly-neutral mutations follows a distribution denoted by *Φ^−^*(S). With the argument similar to Eq.(6), one can show

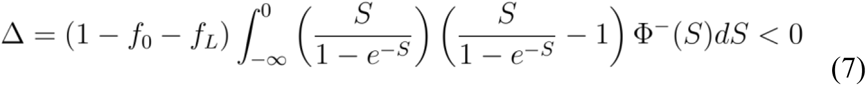

which is always less than 0.

### A new *d*_*N*_/*d*_*S*_ ratio test based on the *Δ* criterion

An interesting question is whether the *Δ* criterion is related to the conventional *d_N_*/*d_S_* test. To this end, we invokes the *H*-measure (Gu et al 1995; Gu 2007), which is defined by

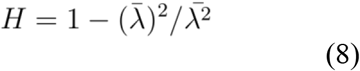

Ranging from 0 to 1, a high value of *H* indicates a high degree of rate variation among sites, and *vice versa*. After some rearrangements of Eq.(8), we have

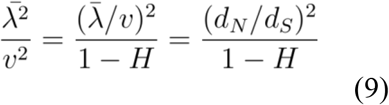

where 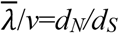 is applied according to Eq.(2). It follows that the relationship between *Δ*, *H* and *d_N_*/*d_S_* is given by

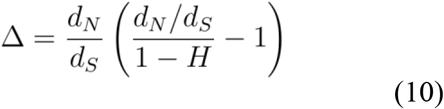

Hence, the connection between *Δ* and *d_N_*/*d_S_* is straightforward: *Δ<0* indicates *d_N_*/*d_S_*<1-*H*, *Δ=0* indicates *d_N_*/*d_S_*=1-*H* and *Δ>0* indicates *d_N_*/*d_S_*>1-*H*, respectively. Accordingly, we formulate a new *d_N_*/*d_S_* ratio test based on the *Δ* criterion (Table 1)

*(i)* The null hypothesis is *d_N_*/*d_S_*=1-*H*, expected by the strictly neutral evolution while some sites are subject to very strong purifying selections (invariable sites).
*(ii)* Rejection of the null hypothesis because of *d_N_*/*d_S_*>1-*H* suggests a positive selection whereas some functionally important sites are virtually invariable.
*(iii)* Rejection of the null hypothesis because of *d_N_*/*d_S_*<1-*H* provides the evidence for the nearly-neutral model instead of the strictly-neutral model with some invariable sites.

**Table 1.**
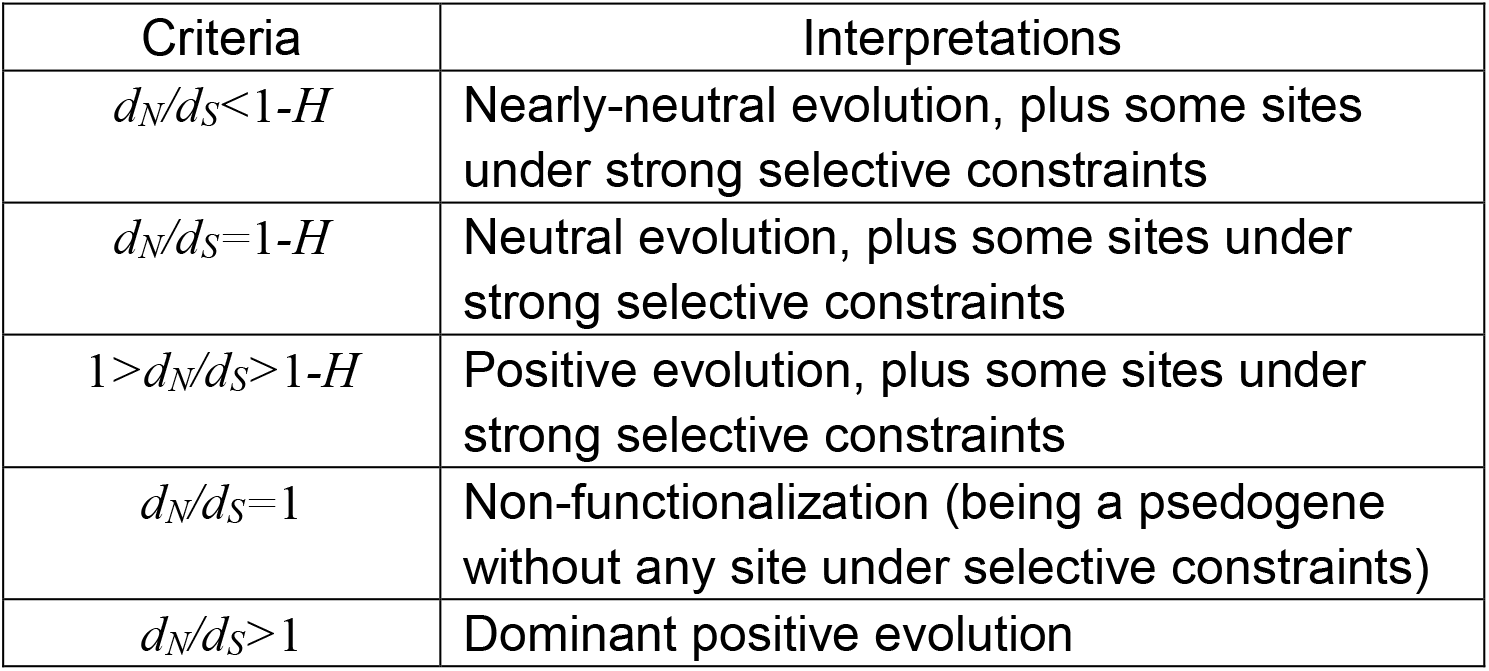
A summary of *Δ*-*d_N_*/*d_S_* analysis

In other words, the new method has two features: (*i*) to detect signal of positive whereas some sites may be subject to very strong constraints (invariable sites); and (*ii*) to distinguish between the nearly neutral model and the strictly-neutral model with invariable sites.

### Calculation of *H*

The remaining question is how one can estimate the parameter *H*, which is fundamental in the new *Δ-d_N_*/*d_S_* test. In the following we discuss types of datasets.

#### Multiple-alignment of protein sequences of a gene

Following Gu (2007), we propose a computational based on a protein sequence phylogeny that can be reliably reconstructed. Let 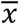 and *V*(*x*) be the mean and variance of number of changes per amino acid site, respectively. Suppose that amino acid changes at a site follow a Poisson process along the phylogeny, whereas the evolutionary rate (λ) varies among sites. Then one may write 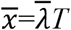 and 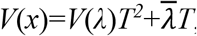, respectively, where *T* is the total evolutionary time along the phylogeny. These two equations together directly lead to an estimate of *H* by

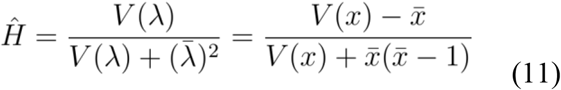

Given the phylogeny, we implement two methods to infer the number of changes per amino acid site: the first one is the parsimony method to infer the minimum-required number of changes per site, and the second one is the method of Gu and Zhang (1997) to estimate the (bias-corrected) number of amino acid changes at each site.

#### Cancer somatic mutations of a gene

The new *Δ-d_N_*/*d_S_* test, which can be applied to study cancer evolution after some specifications, called the *Δ-C_N_*/*C_S_* test. An interesting question is whether the *Δ* criterion is related to the conventional *d_N_*/*d_S_* test. Let *H* be the relative measure for the variation of the rate of somatic mutations among sites. For cancer genomic data, it is straightforward to calculate 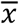 and *V*(*x*), the mean and variance of number of somatic mutations per amino acid site, respectively, which directly lead to an estimate of *H* by Eq.(11).

## Results and Discussion

### Data availability

#### Vertebrate protein sequence data

The protein sequences and homology information of the human and other seven vertebrate genomes (mouse, dog, cow, chicken, Xenopus, fugu, zebrafish) were downloaded from Ensembl EnsMart (http://www.ensembl.org/Multi/martview). Though there are several annotated homology relationships between the human and other genomes in Ensembl, we only considered those pairs of genes annotated as UBRH (Unique Best Reciprocal Hit, meaning that they were unique reciprocal best hits in all-against-all BlastZ searches) to be orthologous. In our study, we used genes that have their orthologs across all eight genomes, i.e. V8 ortholog set. Protein sequences of V8 ortholog set were downloaded from Ensembl; if a gene has multiple protein sequences, usually caused by alternative splicing, only the longest sequence was kept. For each V8 ortholog set, homologous proteins were aligned by the program T-coffee using the default parameters (Notredame et al. 2000). The number of synonymous substitutions per synonymous site (*dS*) and the number of nonsynonymous substitutions per nonsynonymous site (*dN*) between human and mouse orthologs were also retrieved form Ensembl EnsMart, which were estimated by PAML package (Yang 1997) using the likelihood method.

#### Cancer genomics data

We extracted the cancer somatic mutations of 10,224 cancer donors from TCGA PanCanAtlas MC3 project (https://gdc.cancer.gov/about-data/publications/mc3-2017), which includes 3,517,790 somatic mutations in the coding regions, 2,035,693 of which causing amino acid changes in cancers (missense mutations). After filtering out some redundancies according to the criterion of Bailey *et al.*(2018), this left us with 9,078 samples with 1,461,387 somatic mutations in the coding regions, which consisted of 793,577 missense mutations that were used in our study. Sequences and annotations of human genes were extracted from the Ensembl database (http://www.ensembl.org, GRCh37, Release 75). For each gene, we chose the transcript which is most commonly used in TCGA.

Next, genes that have no amino acid site with at least three recurrent missense mutations were excluded because of the lack of degrees of freedom for calculating the *H* value; the final dataset has 4,047 genes. The list of 538 known cancer genes was compiled by merging the list of 253 cancer genes with the missense mutation type from the Cancer Gene Census Tier 1 (COSMIC GRCh37, V89)(Tate et al. 2019), 127 significant mutated genes (SMGs) reported by Kandoth, et al. (2013), 260 SMGs reported by Lawrence, et al. (2013) and 299 cancer driver genes reported by Bailey, et al. (2018).

### Nearly-neutral selection dominates protein evolution in vertebrates

Briefly, the estimation procedure was as follows. (*i*) Infer the phylogenetic tree from a multiple alignment of homologous protein sequences. Although there is no methodological preference, we require that the inferred tree topology should be roughly the same over methods. (*ii*) Estimate the nonsynonymous to synonymous ratio (*d_N_*/*d_S_*) from closely related coding sequences that satisfy *d_N_*/*d_S_* <1. (*iii*). Estimate the *H*-measure for rate variation among sites according to Eq.(11).

We analyzed three independent datasets (Su et al. 2009): (*a*) 342 vertebrate genes from eight species; (*b*) 400 Drosophila genes from 12 species; and (*c*) 380 yeast genes from five species. Impressively, in each dataset, almost all genes satisfy the condition of *d_N_*/*d_S_*<1-*H*, suggesting that nearly-neutral evolution, rather than neutral evolution plus strong negative selection, explains the pattern of protein evolution (**Table 1**). Further, we analyzed 4336 vertebrate genes, while the *d_N_*/*d_S_* ~ (1-*H*) scatter plotting was shown in **Fig.1**. Almost all of genes are below the line of *d_N_*/*d_S_*=1-*H*, or *d_N_*/*d_S_*<1-*H*. We thus conclude that nearly-neutral model is sufficient to explain the genome-wide pattern of protein evolution in vertebrates.

**Fig.1.**
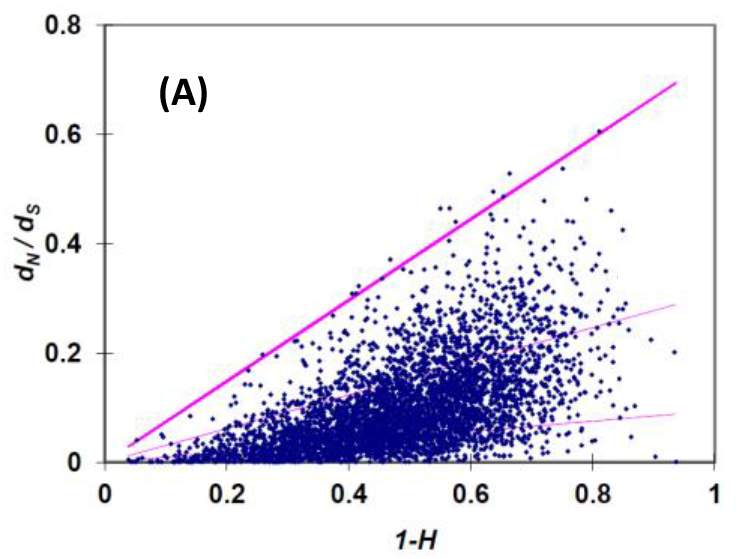
The *d_N_*/*d_S_* ~ (1-*H*) scatter plotting of 433 vertebrate genes, showing that almost all of them are below the line of *d_N_*/*d_S_*=1-*H*, or *d_N_*/*d_S_*<1-*H*.

### Evolution of cancer somatic mutations: Dominant positive selection under strong functional constraints

The theory of clonal evolution in cancer biology claims that cancer cells emerge through random somatic mutations from a single cell and genetically diverge to disparate cell subclones and successive clones in cancer cell replication (Stratton et al. 2009; Vogelstein et al. 2013). Ultimately, cells alter one or few crucial pathways and acquire the hallmarks of cancer through somatic mutations followed by cancer-specific positive selections (Martincorena and Campbell 2015; Bailey et al. 2018). The argument that carcinogenesis is a form of evolution at the level of somatic cells suggests that our understanding of cancer initiation and progression can be benefited by the molecular evolutionary approaches. One well-known example is to estimate the rate ratio of somatic nonsynonymous to synonymous substitutions (*C_N_*/*C_S_*) of a protein-encoding gene in cancers, but resulted in inconsistent conclusions (Dees et al. 2012; Lawrence et al. 2013; Reimand and Bader 2013; Schroeder et al. 2014; Porta-Pardo et al. 2014; Mularoni et al. 2016; Martincorena et al. 2017; Zhou et al. 2017; Weghorn and Sunyaev 2017).

Thanks to the advances in cancer genomics, The Cancer Genome Atlas (TCGA) has accumulated millions cancer somatic mutations from nearly ten thousand tumor-normal pairs. We applied the new *Δ−C_N_*/*C_S_* analysis to explore the major pattern of selection modes in cancer evolution, where *C_N_*/*C_S_* of each gene is calculated by the method of Zhou et al. (2917). First we consider 294 cancer genes. About 70% of them showed *C_N_*/*C_S_* >1, about 30% of them showed 1-*H*<*C_N_*/*C_S_* <1, whereas only one gene showed *C_N_*/*C_S_*<1-*H* (Table 2). Together we propose the following scenario. Driver mutations, i.e., those initiate and facilitate carcinogenesis, confer a selective advantage on cancer cells (positive selection), leading to *C_N_*/*C_S_*>1 for the majority of cancer-related genes. Impressively, up to 30% cancer genes revealed the pattern of mutations under weak positive selection, and mutations under strong purifying selection that were ultimately eliminated from the cancer cell population (Zhou et al. 2017; Weghorn and Sunyaev 2017). Moreover, our analysis suggests that passenger somatic mutations, those unrelated to the process of carcinogenesis, are likely be selectively neutral, rather than nearly neutral due to reduced effective clonal size (Kandoth et al. 2013), because *C_N_*/*C_S_*<1-*H* has been shown highly unlikely. Our analysis in cancer genes has been confirmed by a genome-wide analysis (**Fig.2**).

**Table 2.** Constrting patterns between vertebrate protein sequence evolution and the evolution of cancer somatic mutations.

| Vertebrate protein sequence alignments |  |  |  |
| --- | --- | --- | --- |
| | $d_N/d_S < 1-H$ | $1-H < d_N/d_S < 1$ | $d_N/d_S > 1$ |
|  | 4335 | 1 | 0 |
| Cancer genomic data |  |  |  |
| | $C_N/C_S < 1-H$ | $1-H < C_N/C_S < 1$ | $C_N/C_S > 1$ |
| *Cancer-related* | 1 (0.3%) | 87 (29.6%) | 206 (70.1%) |
| *Genome-wide* | 30 (0.7%) | 2245 (55.5%) | 1772 (43.8%) |

**Fig.2.**
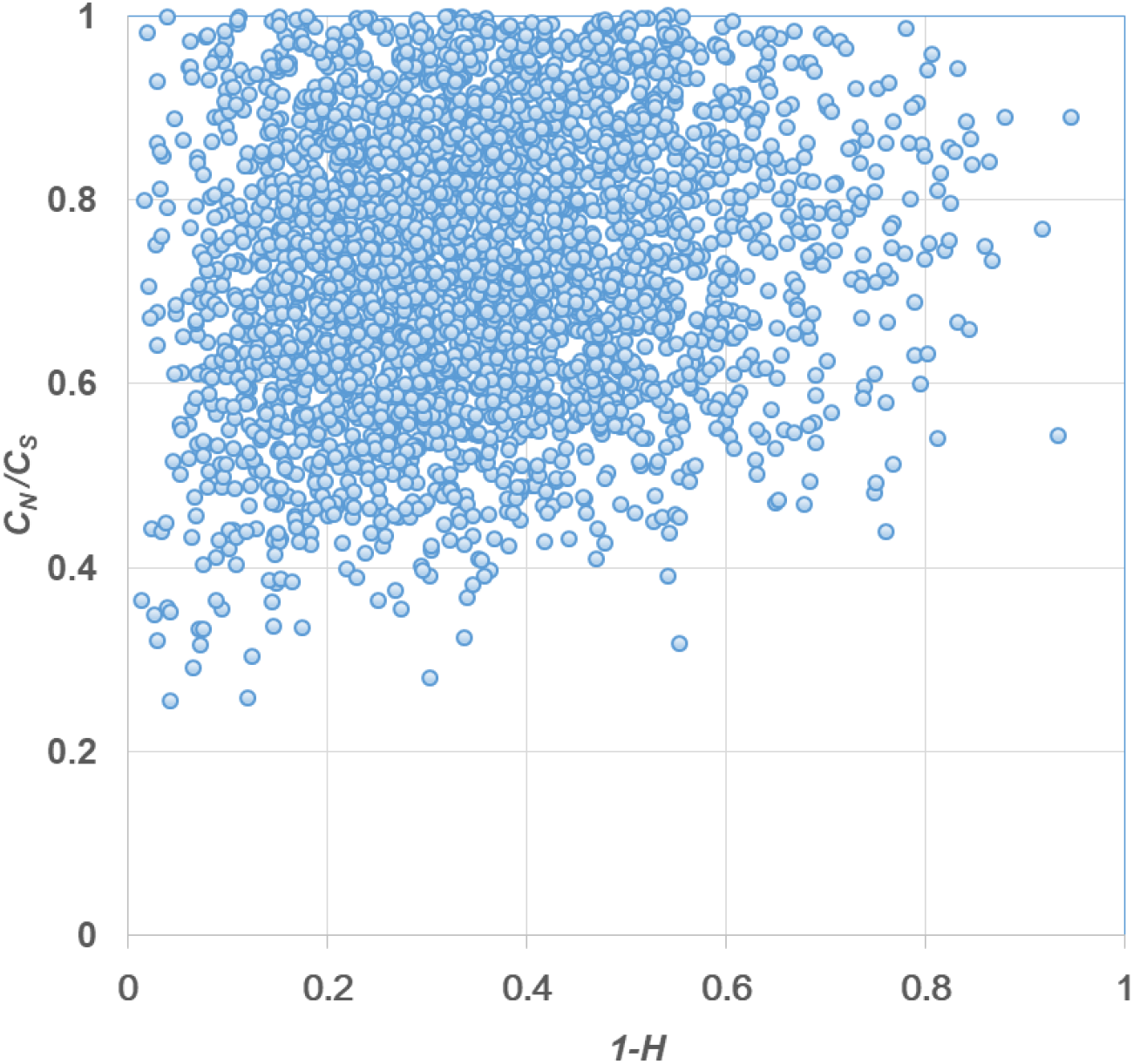
The *C_N_*/*C_S_* ~ (1-*H*) scatter plotting of 2275 human genes with *C_N_*/*C_S_*<1, showing that most of them are above the line of *C_N_*/*C_S_*=1-*H*, or *C_N_*/*C_S_*>1-*H*.

### Further extensions of the *Δ*-*d*_*N*_/*d*_*S*_ test

We anticipate that the new criterion as the null hypothesis of neutrality with strong functional constraints (*d_N_*/*d_S_*=1-*H*) may have many potential applications as it can be implemented into a variety of existed *d_N_*/*d_S_*-related analyses. The challenge is how to estimate the *H* measure accurately when the observed variation of nonsynonymous mutations or substitutions is not statistically sufficient. While it is ideal to estimate *d_N_*/*d_S_* and *H* simultaneously, we found that the mathematical modeling can be sophisticated, leading to computationally difficulty in practice. Technically, one may treat *H* as a known constant in the statistical analysis. In the following we discuss two examples briefly.

#### Δ-d_N_/d_S_ test in the relative rate analysis

Consider the related rate (fast or slow) between two species 1 and 2, given the third species as outgroup. When a relative rate test shows a faster molecular evolution in lineage-1 than lineage-2, the challenge is to determine whether fast-evolution in lineage-1 is driven by adaptive evolution. Let [*d_N_*]_*k*_, [*d_S_*]_*k*_ and [*d_N_*/*d_S_*]_*k*_ be the nonsynonymous distance, synonymous distance and the ratio in the *k*-th lineage, respectively, *k*=1, 2 and 3, respectively. Under the *Δ*-*d_N_*/*d_S_* test, the null hypothesis is that lineage-specific fast-evolution follows a strictly neutral evolution whereas some sites are under strong functional constraints, which predicts [*d_N_*/*d_S_*]_1_=1-*H*, or [*d_N_*]_1_=(1-*H*)[*d_S_*]_1_. Meanwhile, in lineage-2 the *d_N_*/*d_S_* ratio represents the ancestral ratio of the protein sequence conservation, that is, [*d_N_*/*d_S_*]_2_ =[*d_N_*/*d_S_*] in general. Therefore, one can establish the connection between *Δ*-*d_N_*/*d_S_* and the fast-evolving ratio *R_12_*=[*d_N_*]_1_/[*d_N_*]_2_ under the assumption of [*d_S_*]_1_=[*d_S_*]_2_ (synonymous rates are the same between lineages), namely,

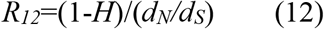

One can develop a statistical procedure to test the null hypothesis of Eq.(12). Rejection of the null hypothesis may indicate a positive selection if [*d_N_*]_1_/[*d_N_*]_2_>*R_12_*, or indicate either functional relaxation or reduced effective population size if [*d_N_*]_1_/[*d_N_*]_2_<*R_12_*. In the case when the two species are not closely related so that the estimation of *d_S_* could be subject to a large sampling variance, one may use the estimate of *d_N_*/*d_S_* from some reliable reference species.

#### The radical/conservative replacement rate ratio

A major variety of *d_N_*/*d_S_*-related analysis is the *D_r_*/*D_c_* ratio test, the ratio of radical replacement rate (*D_r_*) to conservative replacement rate (*D_c_*) (Dagan et al. 2002; Smith 2003; Hanada et al. 2007; Weber et al. 2014; Figuet et al. 2016; Braun 2018; Chen et al. 2019b). The *D_r_*/*D_c_* ratio analysis postulates that, under positive selection more radical amino acid replacements may occur than conservative replacements, and *vice versa* under negative selection. If we view the *D_r_*/*D_c_* ratio as a proxy to the rate-mutation ration, our *Δ*-*d_N_*/*d_S_* analysis can be applied to the *D_r_*/*D_c_* with some technical modifications.

